# Overestimation of sex differences in psychostimulant activity via comparisons of males and females from different behavioral groups

**DOI:** 10.1101/2024.09.26.615282

**Authors:** Anthony M. Tigano, Martin O Job

**Affiliations:** Department of Biomedical Sciences, Cooper Medical School of Rowan University, 401 S Broadway, Camden, New Jersey, USA

**Keywords:** Sex differences, psychostimulant, cocaine, locomotor activity, median split analysis, normal mixtures clustering

## Abstract

**Background:** There are inconsistencies in the observation of sex differences in baseline activity and psychostimulant activity. To address this, we have developed the MISSING (Mapping Intrinsic Sex Similarities as an Integral quality of Normalized Groups) model. MISSING model proposes that sex similarities are observed when we compare similar behavioral groups of males and females, with sex differences occurring when we compare distinct groups of sexes, but this model has not been tested.

**Methods:** To test this model, we identified within-sex groups of Sprague Dawley rats (male n = 22, female n = 23) by conducted normal mixtures clustering of baseline activity, cocaine activity (as distance traveled in cm over 90 min) and cocaine activity normalized-to-baseline activity (NBA) of all subjects. We employed 2-way ANOVA to determine the impact of within-sex heterogeneity on sex differences. We compared our cluster-based method to current median-split approaches.

**Results:** Our new cluster-based method revealed three distinct clusters, each consisting of both males and females. We determined there were no sex differences in any of the variables when males and females from the same clusters were compared. The within-sex clusters for females were not defined by estrous phase. Median split analysis was ineffective in accurately identifying within-sex groups.

**Conclusions:** Our results validate the MISSING model: there are no sex differences in psychostimulant activity except when we compare males and females from different behavioral groups. This has significant implications for how we proceed with research towards understanding the mechanism governing sex differences in psychostimulant activity.

## Introduction

The importance of sex as a biological variable (SABV) (Becker et al., 2016; Beery and Zucker, 2011; Miller et al., 2017; Prendergast et al., 2014; Shansky and Woolley, 2016; Zucker and Beery, 2019) extends to the field of substance use disorders (SUD) research, including as it relates to psychostimulants (Becker, 1999; Becker and Chartoff, 2019; Becker and Hu, 2008; Becker and Koob, 2016). In preclinical models, psychostimulants increase locomotor activity in males and females. It is generally accepted that there are sex differences in psychostimulant-induced locomotor activity with females showing greater responses than males (Cailhol and Mormède, 1999; Chin et al., 2002; Craft and Stratmann, 1996; Festa et al., 2004; Gaines et al., 2022; Gogos et al., 2017; Harrod et al., 2005; Job et al., 2014a; King et al., 2021; Kuhn et al., 2001; Leong et al., 2016; McDougall et al., 2020, 2015; Milesi-Hallé et al., 2007, 2005; Ohia-Nwoko et al., 2017; Parylak et al., 2008; Robison et al., 2017; Schindler et al., 2002; Schindler and Carmona, 2002; Segarra et al., 2010; Sershen et al., 1998; Siegal and Dow-Edwards, 2009; Šlamberová et al., 2013, 2011; Thomsen and Caine, 2011; Van Haaren and Meyer, 1991; Van Swearingen et al., 2013; Wissman et al., 2011; Zhou et al., 2012), *but this is not always the case* (Cailhol and Mormède, 1999; Carroll et al., 2007; Collins et al., 2015; Gaines et al., 2022; Martz et al., 2023; McDougall et al., 2018; Ramos et al., 2020; Risca et al., 2020; Thomsen and Caine, 2011; Zombeck et al., 2010). In addition to sex differences in psychostimulant-induced locomotor activity *per se*, there are also sex differences in baseline locomotor activity (baseline activity) with females showing greater activity than males (Cailhol and Mormède, 1999; Chin et al., 2002; Milesi-Hallé et al., 2007, 2005; Ohia-Nwoko et al., 2017; Ramos et al., 2020; Segarra et al., 2010; Sershen et al., 1998; Simpson et al., 2012; Simpson and Kelly, 2012; Van Haaren and Meyer, 1991; Van Swearingen et al., 2013; Zombeck et al., 2010), *but not always* (Ohia-Nwoko et al., 2017; Robison et al., 2017; Schindler et al., 2002; Schindler and Carmona, 2002). Thus, there can be sex differences in both baseline activity, psychostimulant activity or both.

Baseline activity can regulate/predict subsequent psychostimulant effects, including locomotor activity (Bernardi and Spanagel, 2014; Glick and Milloy, 1973; Irwin et al., 1958; Ohia-Nwoko et al., 2017; Perkins, 1999; Rapport and Dupaul, 1986; Simpson et al., 2012). When we consider that psychostimulant-induced activity is an event that takes place subsequent to (and relative to) baseline activity, psychostimulant-induced activity may be a more accurate variable when expressed as a function of baseline activity. Thus, a combination of baseline activity, psychostimulant activity and another variable that integrates both of these variables (psychostimulant activity normalized-to-baseline activity) may altogether represent a more sensitive group of variables not just for the detection of sex differences but also an identification of within-sex groups. Not all variables are sensitive enough to detect sex differences (Madhuranthakam and Job, 2024), and as such it is important to employ appropriate variables.

Males and females do not represent behaviorally-homogenous populations (Brown et al., 2015; Carreira et al., 2017; Carroll et al., 2007; Davis et al., 2008) and there may be an impact of within-sex heterogeneity on sex differences. When males and females of different behavioral types/groups are compared, it is often observed that there are sex differences or sex similarities, depending on the groups compared. For example, when male and female rats that were selectively-bred for differences in their locomotor response in a novel environment – the so-called High Responder (HR) and Low Responder (LR) rats – were assessed for locomotor activity in a novel environment, it was observed that there were no sex differences when HR males and females were compared, but there were sex differences when LR males and females were compared (Davis et al., 2008). For male and female HR and LR C57BL/6 mice, there were no sex differences with respect to locomotor activity when comparisons were conducted for males and females in the same group (males versus females HR, males versus females LR) (Carreira et al., 2017). Similarly, for rats bred for voluntary running and classified as low voluntary running (LVR) and high voluntary running (HVR) groups, there were no sex differences in locomotor activity when males and females within each group were compared (Brown et al., 2015). Furthermore, high saccharin consuming females showed significantly less locomotion compared to high saccharin consuming males following saline injections but more locomotor activity following the first cocaine injection whereas low saccharin consuming females showed similar locomotion compared to low saccharin consuming males following saline injections and there was no sex differences in locomotor activity after the first cocaine injection (Carroll et al., 2007). For locomotor activity per se and for psychostimulant-induced locomotor activity, sex differences were detected when subjects were handled prior to locomotor activity assessments, but no sex differences were detected for non-handled male and female rats (West and Michael, 1988).

From the above observations, we developed a model which we termed the MISSING (Mapping Intrinsic Sex Similarities as an Integral quality of Normalized Groups) model (see Figure 1B), compared to the current model in Figure 1A. *The idea here is to group similar individuals regardless of sex first, and thereafter determine the impact of group on sex differences (Figure 1B)*. This is a departure from the field where subjects are grouped by sex prior to further assessments (Figure 1A). The MISSING model proposes that 1) there are no sex differences when we compare males and females within the same behavioral group, 2) sex differences occur when we compare males and females from different groups and is tantamount to a group effect and not a biological sex effect *per se*, and 3) even if/when we detect sex differences between males and females in the same behavioral group, these differences will not be as significant as differences between males and females from different groups. The MISSING model has not been tested.

**Figure 1:**
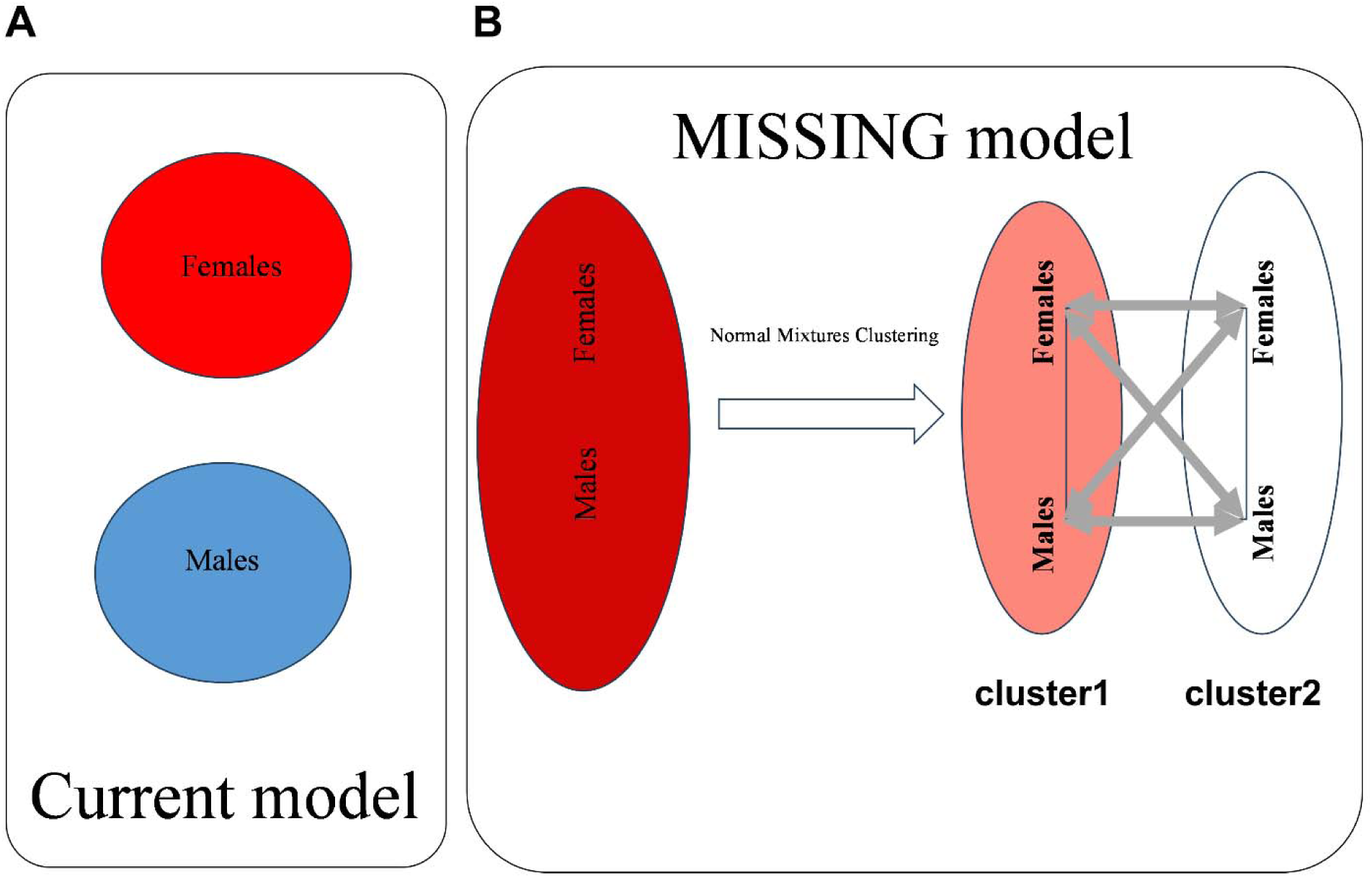
Sex differences as an overestimation of differences between males and females in different, not the same, behavioral groups: the MISSING model. Fig A represents the current model wherein, prior to comparisons, males and females are assumed to represent distinct groups based on biological sex. Fig B is the MISSING (Mapping Intrinsic Sex Similarities as an Integral quality of Normalized Groups) model which does not assume that males and females represent distinct groups prior to comparisons, instead it conducts normal mixtures clustering of all individuals regardless of sex to identify the groups they belong to and afterwards assesses the impact of sex. The square in Fig B captures two clusters (cluster1 and 2) consisting of both males and females. The thick and thin lines around the square represent differences and similarities, respectively. MISSING model proposes that 1) there are no sex differences when we compare males and females within the same behavioral group, 2) sex differences are observed when we compare males and females from different groups, and 3) even if/when we detect sex differences between males and females in the same behavioral group, these differences will not be as significant as differences between males and females from different groups. Sex differences in the current model (Fig A) are likely due to comparisons of males and females from different clusters as in Fig B. The goal of this study was to test this model.

The goal of this study was to validate the MISSING model (Figure 1), including comparing/comparing with current models. To test our model, we designed experiments, in male and female Sprague Dawley rats to first identify groups/clusters from all subjects regardless of sex and thereafter assess the interaction between SEX and cluster. We employed normal mixtures clustering analysis of the following variables: baseline activity, cocaine activity and cocaine activity NBA (cocaine activity normalized-to-baseline activity) for all subjects whether they be male or female *before* we employed 2-way ANOVA to determine the impact of within-sex heterogeneity on sex differences. To identify within-sex groups in the current model we employed the median split analysis of baseline activity and cocaine activity for each sex before we employed 2-way ANOVA to determine if there were SEX × group interactions. Our methods, results and discussions are below.

## Methods

Experiments and animal care were in accordance with the Institute of Animal Care and Use Committee of Emory University and followed the guidelines outlined in the National Institutes of Health (NIH) *Guide for the Care and Use of Laboratory Animals*. We employed a total of forty-five (45) male and female adult Sprague-Dawley rats. These rats were obtained from Charles River Laboratories (Wilmington, MA). They were acclimatized to the housing facility for at least one week before any experimental procedures were conducted. They were provided rat chow and water ad libitum and maintained on a 12-hour light: dark cycle (lights on at 7 am). We conducted all experiments between 10:00 AM and 6:00 PM.

### Assessments of locomotor activity

These were done as in previous studies (Job, 2016; Job et al., 2014b, 2013, 2012; Job and Kuhar, 2017, 2012; Kuhar and Job, 2017). Briefly, we assessed locomotor activity in locomotor chambers (Omnitech Electronics, Columbus OH). These locomotor chambers included transparent Plexiglas walls (dimensions of 40 × 40 × 30 cm) and a photocell cage containing 32 photobeams located 5 cm above the floor (Omnitech Electronics, Columbus OH). The experimental apparatus was such that each photocell cage was connected to a computer equipped with software (Digipro; Omnitech Electronics) to measure locomotor activity. Locomotor activity was measured as distance traveled in centimeters (cm). For these experiments, locomotor activity was obtained for 30 min (baseline activity) after which cocaine (10 mg/kg i.p) was injected and locomotor activity was assessed for an additional 90 min (see Figure 2).

**Figure 2:**
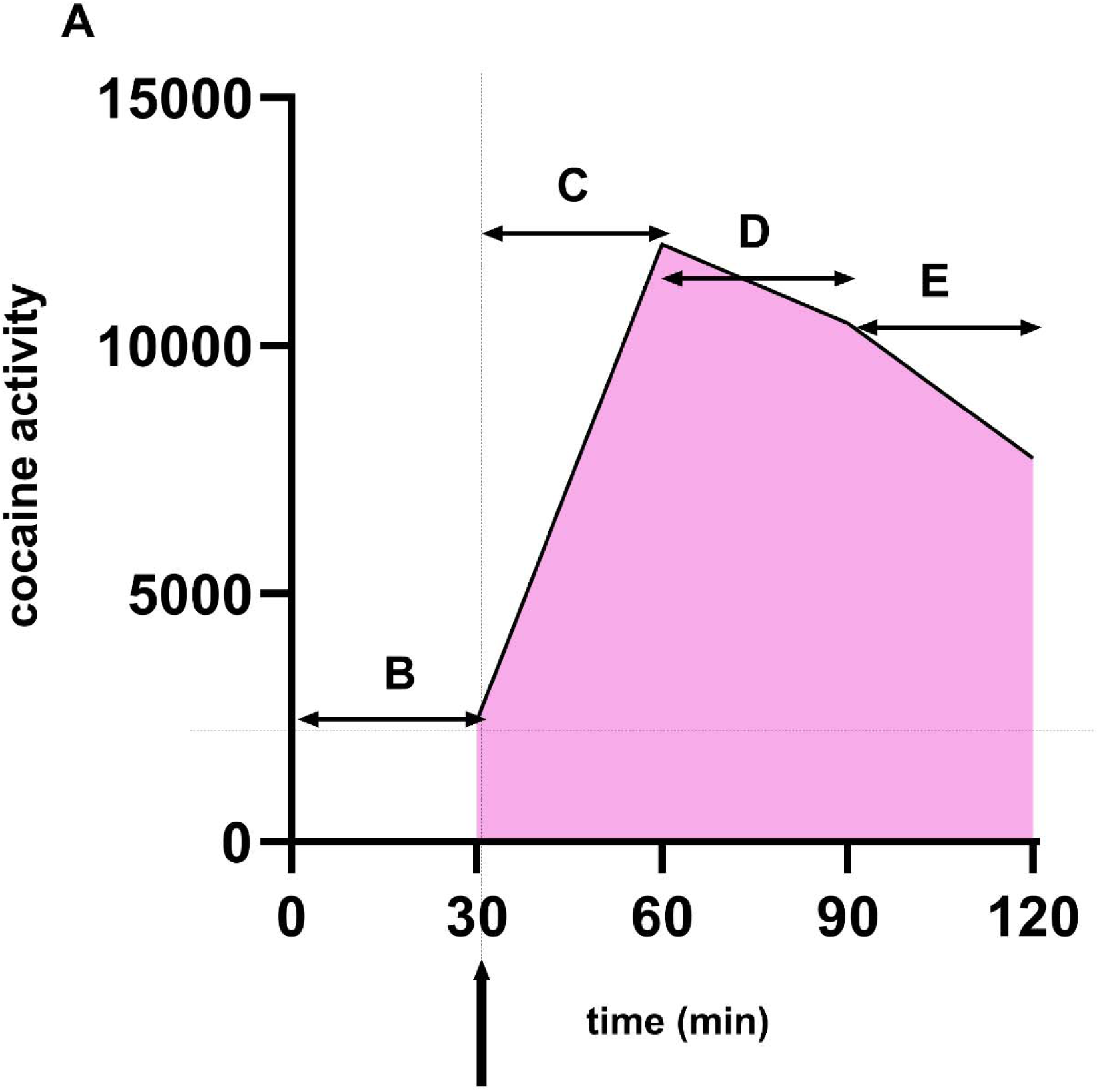
Baseline activity-normalized cocaine activity. The rationale for this variable is that 1. cocaine activity is a change in activity from baseline activity, and 2. while occurring cocaine activity is always relative to baseline activity (not zero activity). Therefore, cocaine activity may be expressed as an activity *continuously* normalized to baseline activity throughout its duration over time. The figure shows how we derived the baseline activity-normalized cocaine activity variable. The x-axis represents time (min), and the y-axis represents the distance traveled in cm (cocaine activity). B represents baseline activity. The arrow at time = 30 min represents the injection point. The shaded region represents the sum over time (90 min) of the distance traveled and corresponds to the area under the curve (total cocaine activity). To obtain baseline activity-normalized cocaine activity, we divided total cocaine activity by baseline activity with baseline activity adjusted for time (C-E). Thus, baseline activity-normalized cocaine activity = cocaine activity (90 min) / baseline activity estimated for 90 min (baseline activity in 30 min × 3).

### Assessment of estrous cycle phase

This was done as described previously. Briefly, on the day of experimentation and after the experiments had been conducted, a subset of female rats (n =15 out of the n = 23) were lightly anesthetized with isoflurane to permit collection of vaginal samples to determine estrous phase. A plastic pipette containing 50 μL of distilled water was used to collect vaginal secretions. To do this, the pipette tip (with distilled water) was inserted gently into the vagina and fluid injected and withdrawn and placed on glass slides. The slides were allowed to dry at room temperature. When dried, the glass slides were stored at −4°C until staining which was done with cresyl violet. The images were scanned at 20X using Aperio and downloaded from Spectrum. The images were not changed or altered in anyway except the scale was adjusted in Adobe Illustrator Encapsulated PostScript (EPS) to fit into a Word document page. The estrous phase is defined by the preponderant cellular types in the vaginal samples. A proestrus sample predominantly contains nucleated epithelial cells whereas an estrus sample predominantly contains anucleated cornified cells (see examples in Figure 5A), for references see (Ekambaram et al., 2017; Marcondes et al., 2002; McLean et al., 2012).

### Variables

Baseline activity was estimated as distance traveled (cm) in 30 min just prior to cocaine injection. Cocaine activity was assessed as distance traveled in 90 min after cocaine injection. Cocaine activity normalized to baseline activity (cocaine activity NBA) was calculated as shown in Figure 2 and as shown in the equation below:

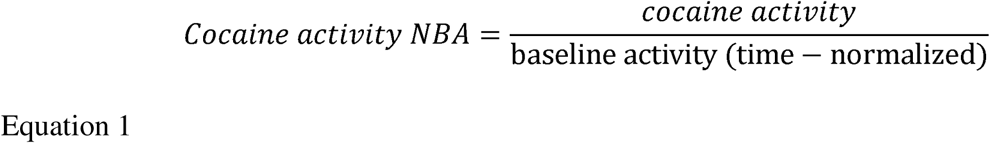

Baseline activity (time normalized) means if cocaine activity is assessed in 90 min and baseline activity is assessed in 30 min, baseline activity (time normalized) would be the baseline activity over the same time of cocaine (90 min) or baseline activity × 3.

### Statistical analysis

Statistical analysis: GraphPad Prism v 9 (GraphPad Software, San Diego, CA), SigmaPlot 14.5 (Systat Software Inc., San Jose, CA) and JMP Pro v 17 (SAS Institute Inc., Cary, NC) were employed for statistical analysis. Grubb’s test was used to determine if there were any significant outliers. Data were expressed as mean ± SEM. For the new model, we employed normal clustering of several variables (baseline activity, cocaine activity and cocaine activity NBA) for every individual, regardless of sex, to identify distinct clusters/groups. We confirmed that these clusters/groups were indeed distinct using One-way ANOVA for variable comparisons and linear regression analysis for comparisons of relationships between variables. Thereafter, we employed Two-way ANOVA to determine if there were main effects of SEX, cluster and SEX × cluster interaction. We compared this new model with the current model. For group identification in the current model, we employed median split analysis of two of the variables (baseline activity, cocaine activity). Statistical significance was set at p < 0.05 for all analyses with Tukey’s post hoc test employed when significance was detected.

## Results

### Sex differences in cocaine activity but not baseline activity or cocaine activity NBA

The mean ± SEM for baseline activity for males (n = 22) and females (n = 23) were 2427 ± 319 and 2874 ± 325 cm, respectively. With regards to baseline activity, Grubb’s test revealed no significant outliers for males or females or all. Unpaired t-tests revealed no significant difference between sexes with regards to baseline activity (P = 0.3314, Figure 3A). Analysis of the data distribution suggests two (2) populations (normally distributed) of males (Figure 3B) and one (1) population (normally distributed) of females (Figure 3C).

**Figure 3:**
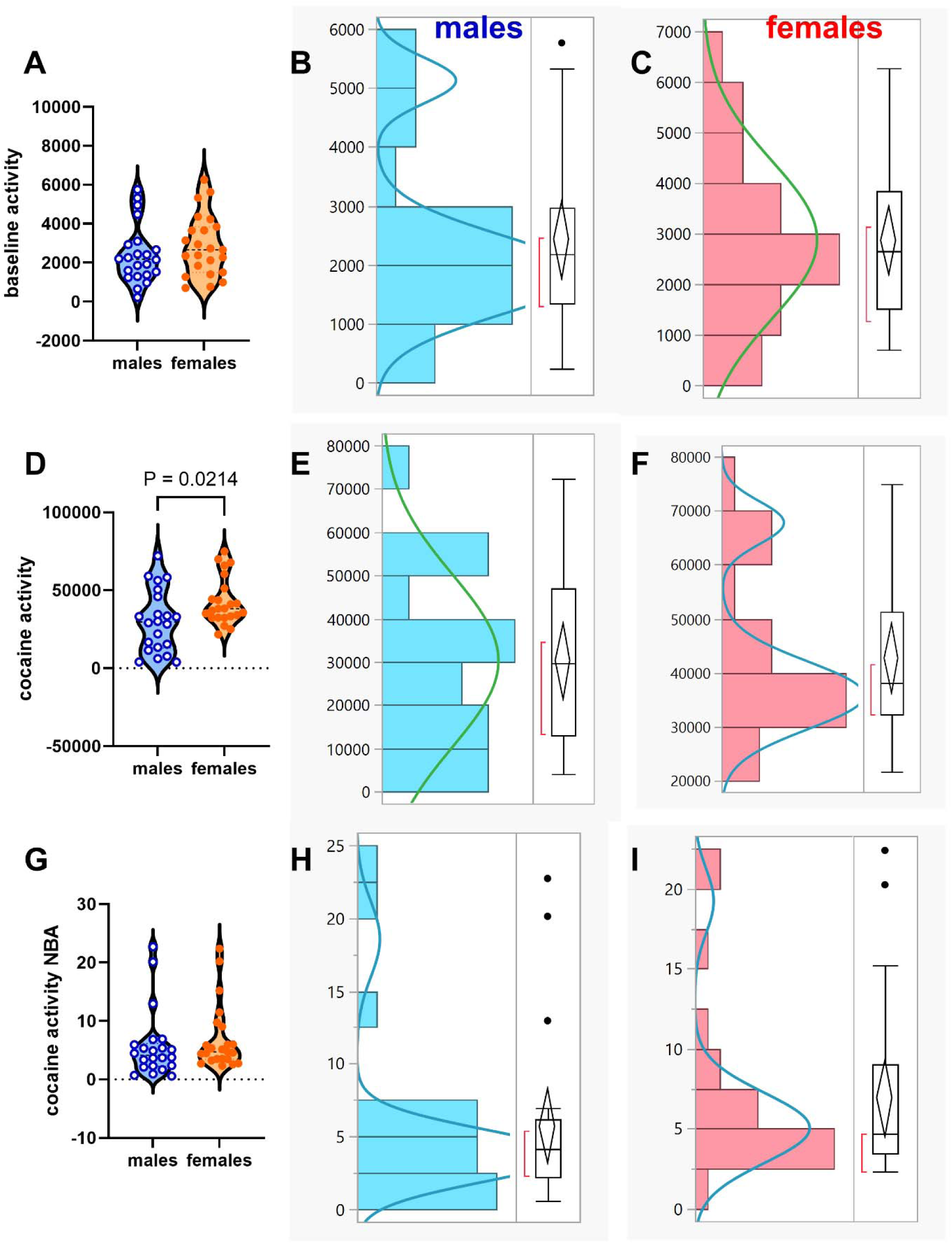
Sex differences in cocaine activity but not baseline activity-normalized cocaine activity: more than one behavioral group of subjects in sample: We employed unpaired t-tests to determine if there were sex differences in 1. baseline activity (Fig A), 2. cocaine activity (Fig D) and 3. cocaine activity NBA (Fig G). There were no sex differences in baseline activity (P = 0.3314, Fig A). We analyzed the distribution for baseline activity values for males (Fig B) and females (Fig C) detecting a mixture of two normally distributed populations for males and one normally distributed population of females. There were sex differences, as expected, in cocaine activity (P < 0.05, Fig D). We analyzed the distribution for cocaine activity for males and females (Fig E-F), and this time it was for the females that a mixture of two populations (normal distribution) was revealed while the males appeared to consist of one population (normally distributed). Interestingly, there was no sex difference in cocaine activity NBA (P = 0.4871, Fig G). Interestingly, analysis of the distribution of cocaine activity NBA revealed mixtures of two populations of male and female subjects (Fig H-I). Grubb’s outlier test revealed an outlier for male and female cocaine activity NBA when analyzed separately, but when males and females were combined Grubb’s test detected no outliers. This suggests that the seeming ‘outliers’ belonged to a different population.

The mean ± SEM for cocaine activity for males (n = 22) and females (n = 23) were 30225 ± 4244 and 42756 ± 3142 cm, respectively. With regards to cocaine activity, Grubb’s test revealed no significant outliers for males, females or males plus females. Unpaired t-tests revealed a significant difference between sexes with regards to cocaine activity (P = 0.0214, Figure 3D). Analysis of the distribution suggests one (1) normally distributed population of males (Figure 3E) and two (2) populations of females (normally distributed) (Figure 3F).

The mean ± SEM for cocaine activity NBA for males (n = 22) and females (n = 23) were 5.73 ± 1.23 and 6.91 ± 1.15, respectively. With regards to cocaine activity, Grubb’s test revealed significant outliers for males and for females but not for all. Unpaired t-tests revealed no significant difference between sexes with regards to cocaine activity NBA (P = 0.4871, Figure 3G). Analysis of the distribution suggests two (2) normally distributed populations of males (Figure 3H) and females (Figure 3I).

### Normal mixtures clustering of baseline activity, cocaine activity and cocaine activity NBA

Normal mixtures clustering of baseline activity, cocaine activity and cocaine activity NBA revealed 3 clusters with all clusters consisting of males and females (Figure 4A-B). We labeled these as cluster1 (n = 9: males n = 3, females n = 6), cluster2 (n = 14: males n = 7, females n = 7) and cluster3 (n = 22: males n = 12, females n = 10). The baseline activity for cluster1 (n = 9), cluster2 (n =14) and cluster3 (n = 22) were 1126 ± 202, 2013 ± 161 and 3691 ± 310 cm, respectively. The cocaine activity for cluster1, 2 and 3 were 47058 ± 5458, 31985 ± 2187 and 35319 ± 4823 cm, respectively. The cocaine activity NBA for cluster1, 2 and 3 were 15.99 ± 1.82, 5.45 ± 0.22 and 2.94 ± 0.28, respectively. One-way ANOVA revealed significant differences between clusters for baseline activity (F 2, 24 = 20.34, P < 0.0001, Figure 4C) and cocaine activity NBA (F 2, 24 = 82.36, P < 0.0001, Figure 4E), but not for cocaine activity (F 2, 24 = 2.012, P = 0.1464, Figure 4D). The slopes of the relationship between baseline activity and cocaine activity (Figure 4F) were significant for all clusters: cluster1 (F 1, 7 = 7.064, P = 0.0326, R^2^ = 0.50, slope = 19.14 ± 7.203, y-intercept = 25503 ± 9095), cluster2 (F 1, 12 = 34.34, P < 0.0001, R^2^ = 0.74, slope = 11.72 ± 2.00, y-intercept = 8396 ± 4189) and cluster3 (F 1, 20 = 47.76, P < 0.0001, R^2^ = 0.71, slope = 13.06 ± 1.89, y-intercept = -12878 ± 7473), and there were no differences between clusters with respect to slopes (F 2, 39 = 0.5006, P = 0.6100), but there were significant differences with respect to y-intercepts (F 2, 41 = 33.32, P < 0.0001). Similarly (Figure 4G), the slopes of the relationship between baseline activity and cocaine activity NBA were significant for all clusters: cluster1 (F 1, 7 = 11.48, P = 0.0116, R^2^ = 0.62, slope = - 0.007086 ± 0.002092), cluster2 (F 1, 12 = 11.10, P = 0.0060, R^2^ = 0.48, slope = -0.0009442 ± 0.0002834) and cluster3 (F 1, 20 = 7.497, P = 0.0127, R^2^ = 0.27, slope = +0.0004630 ± 0.0001691), and there were significant differences when we compared slopes for clusters 1-3 (F 2, 39 = 26.15, P < 0.0001).

**Figure 4:**
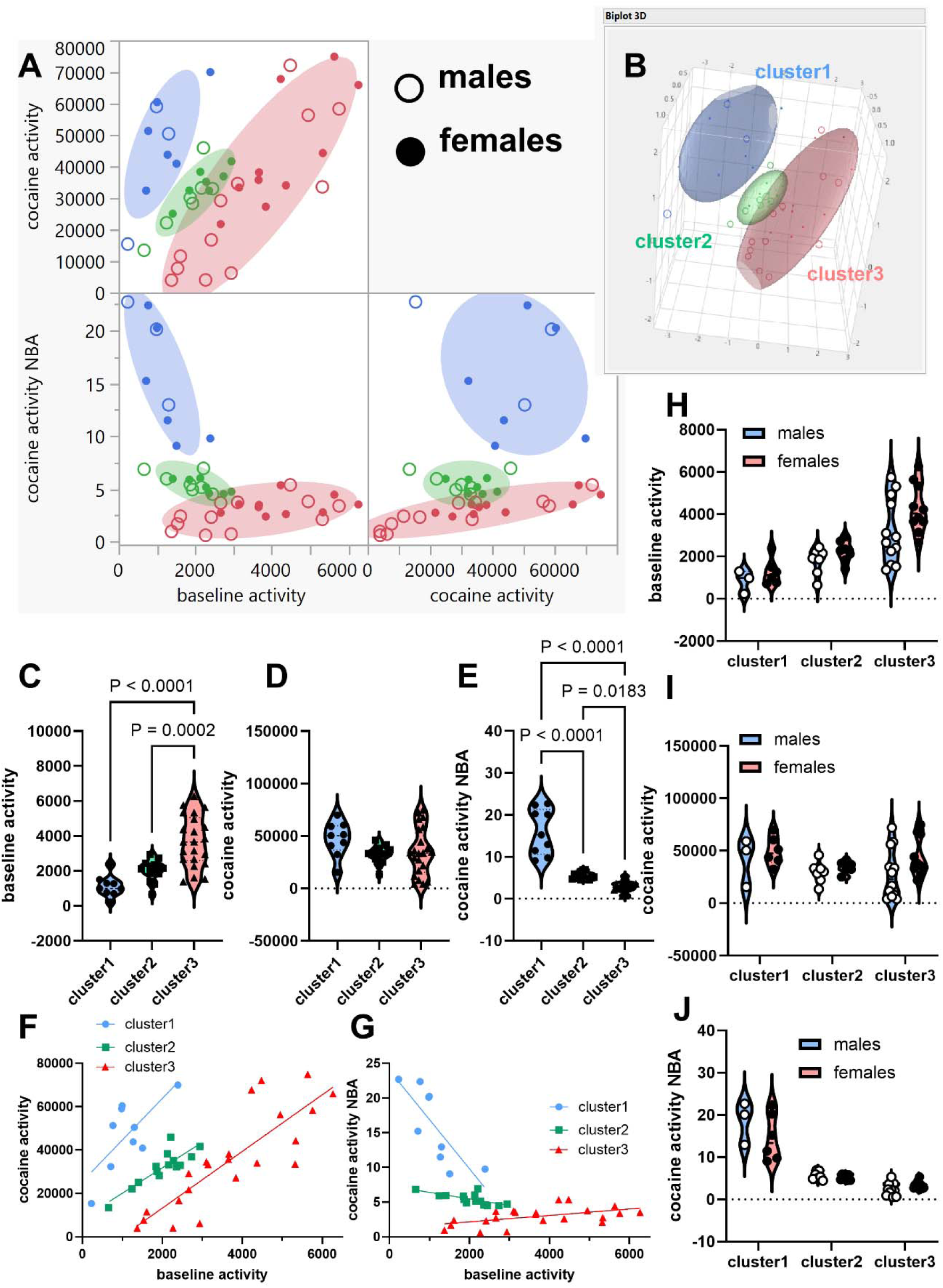
Normal mixtures clustering analysis of baseline activity, cocaine activity and cocaine activity NBA yielded three distinct groups of males and females. We conducted normal mixtures clustering of baseline activity, cocaine activity and cocaine activity NBA for all subjects (n = 45). This analysis yielded three clusters (Fig A-B): cluster 1 (n = 9: males n = 3, females n = 6), cluster2 (n = 14: males n = 7, females n = 7) and cluster3 (n = 22: males n = 12, females n = 10). One-way ANOVA revealed significant differences between clusters for baseline activity (P < 0.0001, Fig C) and cocaine activity NBA (P < 0.0001, Fig E), but not for cocaine activity (P = 0.1464, Fig D). Linear regression revealed a significant relationship between baseline activity and cocaine activity for all 3 clusters (P < 0.05, Fig F), with no differences between clusters with respect to slopes (P = 0.6100), but significant differences with respect to y-intercepts (P < 0.0001). Similarly (Fig G), the slopes of the relationship between baseline activity and cocaine activity NBA were significant for all clusters (P < 0.05) with significant differences when we compared slopes for clusters 1-3 (P < 0.0001). We conducted Two-way ANOVA with SEX (males, females) and clusters (cluster1, cluster2, cluster3) to determine if there was a SEX × cluster interaction and main effects of SEX and cluster. There was no SEX × cluster interaction and main effect of SEX (P > 0.05) for baseline activity (Fig H), cocaine activity (Fig I) and cocaine activity NBA (Fig J). Interestingly, there was a main effect of cluster (P < 0.0001) for baseline activity and cocaine activity NBA, but not for cocaine activity.

Two-way ANOVA with factors SEX (males, females) and clusters (cluster1-3) and dependent variable = baseline activity did not reveal a SEX × cluster interaction (F 2, 39 = 0.4764, P = 0.6244) or a main effect of SEX (F 1, 39 = 3.512, P = 0.0684) but did reveal a main effect of cluster (F 2, 39 = 22.70, P < 0.0001), see Figure 4H. For cocaine activity, Two-way ANOVA did not reveal a SEX × cluster interaction (F 2, 39 = 0.4833, P = 0.6204) or any main effects of SEX (F 1, 39 = 2.948, P = 0.0939) and cluster (F 2, 39 = 1.562, P = 0.2225), see Figure 4I. For cocaine activity NBA, Two-way ANOVA did not reveal a SEX × cluster interaction (F 2, 39 = 2.605, P = 0.0867) or a main effect of SEX (F 1, 39 = 1.957, P = 0.1697) but did reveal a main effect of cluster (F 2, 39 = 87.50, P < 0.0001), see Figure 4J.

Note that there were no sex differences within any cluster but there were sex differences when we compared males and females from different clusters. Sex differences appear to be driven by cluster differences not biological sex *per se*.

### The within-sex clusters for females were not defined by estrous phase

For the females used in the experiments, n = 15 out of the n = 23 were assessed for estrous phase on the day of the experiments (see Methods section, see example in Figure 5A). Of these n = 15 subjects assessed for estrous phase, Cluster 1, 2 and 3 contained n = 5 subjects each (triangles in Figure 5B). Cluster 1 had 1 subject in proestrus and 4 subjects in estrus (Figure 5C, cluster1). Cluster 2 had 4 subjects in proestrus and 1 subject in estrus (Figure 5C, cluster2). Cluster 3 had 2 subjects in proestrus and 3 subjects in estrus (Figure 5C, cluster3).

**Figure 5.**
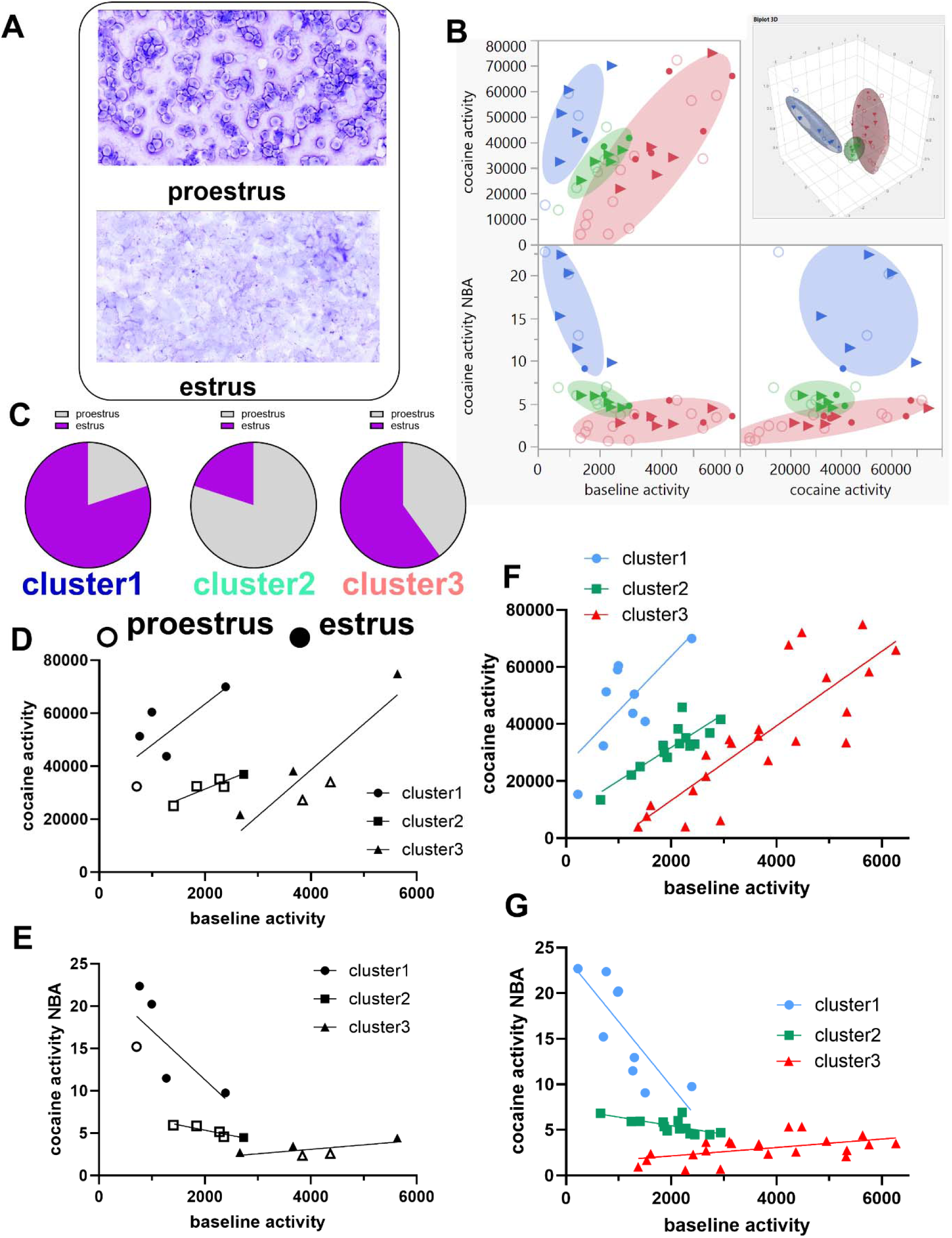
Clusters include mixtures of different estrus stages, but this does not appear to determine the characteristics of the clusters. Fig A shows examples of estrus phases (proestrus and estrus) staged using vaginal lavage and cresol violet stain and cell types observed under the microscope. Fig B shows the females that were assessed for estrus phase (n = 15 with n = 5 subjects in each cluster, see triangles in Fig B) relative to all subjects (n = 45, this is also Figure 4A-B). Fig C shows the relative % for proestrus and estrus in the clusters. Fig D shows the relationship between baseline activity and cocaine activity for these n = 15 females in the three clusters (open shapes represent proestrus and closed shapes represents estrus). Fig E shows the relationship between baseline activity and cocaine activity NBA for these n = 15 females in the three clusters (open shapes represent proestrus and closed shapes represents estrus). Fig F represents the relationship between baseline activity and cocaine activity for all three clusters and for all subjects. Fig G represents the relationship between baseline activity and cocaine activity NBA for all three clusters and for all subjects. The relationship between baseline activity and cocaine activity or cocaine activity NBA for all subjects (male and female) were not different (P > 0.05) when we account for different group compositions of females in estrus versus proestrus, suggesting that the clusters were not determined by estrous cycle phase.

Regardless of group composition (mostly estrus, mostly proestrus or a mixture of both for cluters1-3, Figure 5C), the slopes representing the relationship between baseline activity and 1) cocaine activity (Figure 5D) and/or 2) cocaine activity NBA (Figure 5E) for these females were not different (P > 0.05) from that of the whole cluster that they belonged to (males plus females) (Figure 5D versus 5F, Figure 5E versus 5G). Thus, the differences between females in clusters1-3 cannot be explained by proestrus or estrus phases. Interestingly, females in proestrus tended to have similar baseline activity but lower cocaine activity relative to those in estrus (Figure 5D). Interestingly also, females in proestrus and estrus tended to have similar cocaine activity NBA regardless of cluster.

### The groups identified by the clustering analysis were not identified via median split analysis

Current models typically identify within-sex groups using median split of cocaine activity variables to yield 2 groups – one group above the median termed the high responder group and one group below the median termed the low responder group. Our normal mixtures clustering analysis identified 3 groups (clusters1-3, Figure 4). Median split does not typically identify three groups. This implies that median split analysis would not be effective in identifying the groups we identified.

We employed median split of cocaine activity for males (Figure 6A) and females (Figure 6B). With factors as SEX (males, females) and groups (low cocaine and high cocaine), Two-way ANOVA revealed no SEX × group interaction (F 1, 41 = 2.246, P = 0.1416), but did reveal a main effect of SEX (F 1, 41 = 13.06, P = 0.0008) and group (F 1, 41 = 63.68, P < 0.0001), see Figure 6C. There were n = 22 subjects (n = 11 males and n = 11 females) in the low cocaine groups while there were n = 23 subjects (n = 11 males and n = 12 females) in the high cocaine group, compare with cluster1, 2 and 3 containing n = 9, 14 and 22 subjects (Figure 4).

**Figure 6:**
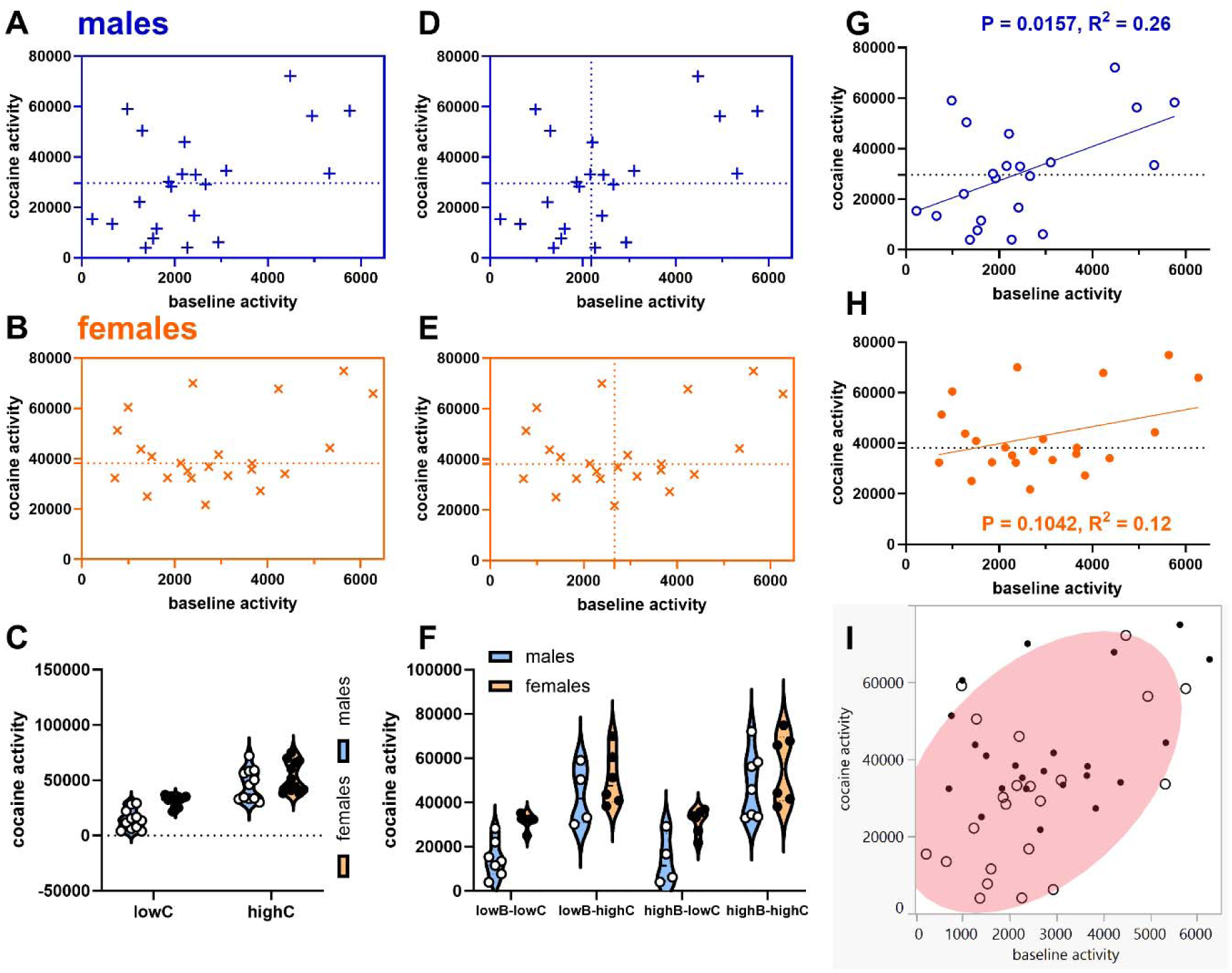
Median split analysis of cocaine activity and/or both baseline activity and cocaine activity yielded groups of males and females that were not the same as the groups obtained from normal mixtures clustering of baseline activity, cocaine activity and cocaine activity NBA. We employed median split of cocaine activity for males (Fig A) and females (Fig B) to realize low (lowC) and high (highC) cocaine activity groups for both sexes. Two-way ANOVA revealed no SEX × group interaction (P = 0.1416) but did reveal a main effect of SEX (P = 0.0008) and group (P < 0.0001). There were n = 22 subjects (n = 11 males and n = 11 females) in the low cocaine groups while there were n = 23 subjects (n = 11 males and n = 12 females) in the high cocaine group, compare with cluster1, 2 and 3 containing n = 9, 14 and 22 subjects (Figure 4). We also conducted median split of both baseline activity and cocaine activity for males and females (Fig D and E) to yield 4 groups (low baseline – low cocaine, low baseline-high cocaine, high baseline-low cocaine and high baseline-high cocaine groups; lowB-lowC, lowB-highC, highB-lowC and highB-highC). 2 × 4 ANOVA revealed no SEX × group interaction (P = 0.1416) but did reveal a main effect of SEX (P = 0.0012) and group (P < 0.0001). However, for males, there was a significant relationship between baseline activity and cocaine activity (P = 0.0157, R^2^ = 0.26, Fig G) but not so for females (P = 0.1042, R^2^ = 0.12, Fig H), with no differences between males and females for these relationship(s) (P = 0.2955). Normal mixtures clustering of baseline activity and cocaine activity all subjects, whether they be male or female, yielded only one cluster (Fig I). In summary, median split analysis does not identify the same groups we identified using normal mixtures clustering, see Figure 4. Also, based on the variables, there was only one cluster of males and females and as such median split analysis overestimates groups (Fig A, B, D and E).

We also conducted median split of both baseline activity and cocaine activity for males and females (Figure 6D-E) to yield 4 groups (low baseline – low cocaine, low baseline-high cocaine, high baseline-low cocaine and high baseline-high cocaine groups). 2 × 4 ANOVA revealed no SEX × group interaction (F 3, 37 = 0.5996, P = 0.1416), but did reveal a main effect of SEX (F 1, 37 = 12.24, P = 0.0012) and group (F 3, 37 = 19.46, P < 0.0001), see Figure 6F. There were n = 7 males and n = 5 females for low baseline – low cocaine, n = 4 males and n = 6 females for low baseline-high cocaine, n = 4 males and n = 6 females for high baseline-low cocaine and n = 7 males and n = 6 females for high baseline-high cocaine groups. Compare these group consistencies with cluster1 (males n = 3, females n = 6), cluster2 (males n = 7, females n = 7) and cluster3 (males n = 12, females n = 10) (Figure 4).

## Median-split and normal mixtures clustering of baseline activity and cocaine activity do not yield the same groups

Linear regression analysis reveals that there is a significant relationship between baseline activity and cocaine activity for males (F 1, 20 = 6.963, P = 0.0157, R^2^ = 0.26, slope = 6.769 ± 2.565, Figure 6G) but not for females (F 1, 21 = 2.885, P = 0.1042, R^2^ = 0.12, slope = 3.364 ± 1.981, Figure 6H). There are no differences between males and females for these relationship(s) (F 1, 41 = 1.123, P = 0.2955). Normal mixtures clustering of baseline activity and cocaine activity all subjects, whether they be male or female, yielded only *one cluster* (Figure 6I). Median split of baseline activity and cocaine activity yielded *4 clusters* (Figure 6D-E).

## Discussion

To explain inconsistencies in the observations of sex differences in baseline and psychostimulant activity, we developed a MISSING model (Figure 1) which proposes that there exists distinct clusters consisting of males and females, with sex similarities within clusters and sex differences between clusters (this sex difference is due to behavioral differences and not biological sex).

Because psychostimulant activity is dependent on baseline activity and is also relative to baseline activity, we utilized three variables – baseline activity, cocaine activity and cocaine activity NBA (Figure 2) to increase our sensitivity of detecting sex differences and within-sex groups, if any. When we do not account for within-sex groups, there are sex differences in cocaine activity, but not in baseline activity and cocaine activity NBA (Figure 3). When we account for within-sex groups, there are no sex differences in baseline activity, cocaine activity or cocaine activity NBA (Figure 4), only cluster differences for baseline activity and cocaine activity NBA. Because there were distinct groups of males and females in our sample, the sex differences observed for cocaine activity (Figure 3B) are likely driven by group differences not biological sex. Indeed, if we compared males and females from different clusters, we would detect sex differences, but these differences are merely group differences, not differences due to biological sex. Our data validates the MISSING model.

We observed that the distinct clusters of female subjects could not be explained by the estrous phase (Figure 5). Our observation the distinct clusters of female subjects could not be explained by the estrous phase (Figure 5) is corroborated by a previous study showing estrous phase could not explain the differential locomotor activity of HR-bred and LR-bred groups of rats – the group-related differences were not due to estrus phase (Davis et al., 2008).

The distinct groups identified using normal mixtures clustering of baseline activity, cocaine activity and cocaine activity NBA did not differ with regards to cocaine activity (Figure 4). The idea for median split is to separate subjects into groups based on high versus low cocaine activity or drug-related activity or novelty-related activity (Merritt and Bachtell, 2013; Nelson et al., 2009; O’Connor et al., 2022). Because median split is applied to categorize subjects based on differential cocaine activity, as such, it would not be sensitive enough to identify distinct groups that have the same cocaine activity (Figure 4). Further, the groups identified by median split of two current variables (Figure 6) were not of the same group consistency we identified using normal mixtures clustering analysis of the same variables and a new normalized variable (Figure 4). But even when we focus only on baseline activity and cocaine activity, median split yielded different groups (high versus low responders, Figure 6A, B, D, E) while normal clustering analysis of the same variables yielded only one group (Figure 6I). We have already published a report showing that median split analysis is ineffective in identifying consistent groups when several variables are considered (Castaneda and Job, 2024). Here, we again highlight the problems with this analytical procedure.

We showed that baseline activity was positively correlated to psychostimulant-induced activity for males (Figure 6G) but not females (Figure 6H). This is similar to reports showing that baseline activity was positively correlated to methamphetamine-induced activity in male but not female BALB/c mice (Ohia-Nwoko et al., 2017), though we also show no sex differences with regards to this correlation (baseline activity versus cocaine activity).

In summary, we determined that when we conducted normal mixtures clustering of all data, regardless of sex, we identified clusters consisting of males and females. We determined that there were sex similarities in behavior when we compared males and females within a cluster. We observed sex differences when we compared males from one cluster with females from another cluster, but this assumed sex difference is not due to biological sex per se but akin to the cluster difference in behavior(s). We noted the limitation of the current model which groups subjects by sex before any analysis. We noted (another) limitation of the median split approach.

This work has significant implications for how we proceed with research towards understanding the mechanism governing sex differences in psychostimulant activity. Without applying the MISSING model, the analysis of molecular mechanisms for sex differences may actually be mostly identifying group differences, not biological sex differences. To understand sex differences, we must either 1) focus on males and females within the same cluster or 2) account for the impact of group differences on sex differences if we compare males and females from different clusters. This work will advance the field of sex differences research.

## Acknowledgements

The authors wish to acknowledge Michael J Kuhar in whose laboratory MOJ conducted the behavioral experiments. All authors contributed to data analysis and to the writing of the manuscript. MOJ designed and conducted the behavioral experiments and statistical analysis. The authors acknowledge the support of NIH grants DA15162 and DA015040. Also this work was funded by the National Center of Research Resources P51RR165 and was supported by the Office of Research Infrastructure Programs/OD P51OD11132. This work was also supported by the Francis Lax Fund for Faculty Development at Rowan University. This work was also supported by startup funds from Rowan University, Camden, New Jersey.

